# ESBL plasmid compatibility with the surrounding microbial community influences ESBL gene survival under CRISPR-antimicrobial targeting

**DOI:** 10.1101/2025.04.15.648872

**Authors:** Cindy J. Given, Ilmur Jonsdottir, Krista Norvasuo, Petra Paananen, Pilvi Ruotsalainen, Teppo Hiltunen, Marianne Gunell, Antti J. Hakanen, Matti Jalasvuori, Reetta Penttinen

**Affiliations:** University of Jyväskylä, Department of Biological and Environmental Science, Nanoscience Center, Survontie 9C, Jyväskylä, Finland; Department of Biology, FI-20014 University of Turku, Turku, Finland; Department of Clinical Microbiology, Turku University Hospital and University of Turku, Turku, Finland

**Author notes:** Corresponding author: Reetta Penttinen.

**Keywords:** Conjugative CRISPR-antimicrobial, ESBL, multispecies evolution experiment

## Abstract

Bacteria often acquire resistance against antibiotics through the transfer of conjugative resistance plasmids. Hence, it is vital to develop strategies to mitigate the dispersal of antimicrobial resistance (AMR). CRISPR-based antimicrobial tools offer a sequence-specific solution to diminish and restrict the dissemination of antimicrobial resistance genes among bacteria. CRICON (CRISPR via conjugation) is an antimicrobial CRISPR tool that has been shown to efficiently reduce multi-resistance when targeting ESBL (Extended Spectrum Beta-Lactamase) harboring plasmids. However, conjugatively delivered genetic elements may be subjected to bacterial defense, lead to resistance development, and revert the efficiency of the CRISPR tools. Here, we studied the evolutionary consequences of four ESBL-harboring *Escherichia coli* strains targeted by CRICON in a 10-day multispecies microcosm experiment. We show that CRICON reduces the ESBL prevalence within the bacterial community, while the final ESBL persistence depends on the initial community composition. We observed an unexpected survival strategy of an ESBL-plasmid by escaping into a more competitive host species. Further, we show the development of partial resistance against the CRISPR-antimicrobials during the experiment. Our results underline the importance of the ecological and evolutionary factors in multispecies bacterial communities, as they may disrupt the effective use of CRISPR-based antimicrobial strategies via undesired outcomes of targeted therapies against plasmid-bearing multi-resistant bacteria.

## Introduction

Bacterial antimicrobial resistance (AMR) poses a significant global threat to public health. Clinical treatments primarily rely on antibiotic compounds, but bacteria can accumulate antimicrobial resistance genes (ARGs) through conjugative plasmids, leading to the development of multi-resistant bacterial strains. Therefore, it becomes crucial to prolong the use of existing antibiotics, e.g., through applications restricting the flow of AMR plasmids within the microbial communities and/or targeting the antimicrobial resistance genes themselves. Native CRISPR (<u>C</u>lustered <u>r</u>egularly interspaced <u>s</u>hort <u>p</u>alindromic <u>r</u>epeats)-Cas (<u>C</u>RISPR-<u>as</u>sociated) systems are bacterial and archaeal immune systems that facilitate protection against bacteriophages and other invading DNA elements, namely conjugative plasmids. The CRISPR defense mechanism allows bacteria to learn to recognize invading genetic elements in a sequence-specific manner in a process called CRISPR immunization (1,2). In general, CRISPR-based immunity utilizes Cas genes in collaboration with CRISPR spacer arrays that serve as a genetic memorization library to govern resistance against potential re-infections of the intruding phages and plasmids. CRISPR spacers correspond to sequences of the targeted nucleic acid. When processed into gRNAs (guide RNA), they direct the CRISPR defense against the target DNA/RNA via complementary pairing between the gRNA and its target strand (3–5). Binding to the target sequence is mediated by a complex consisting of Cas protein(s) and the gRNA. Depending on the type of the CRISPR-Cas system, Cas proteins with endonuclease activity mediate the subsequent cutting of the target sequence (1,2). With regards to the sequence-specific targeting, CRISPR-based genetic tools allow a programmable approach to target specific genes precisely and thereby diminish AMR genes from bacterial populations (6).

There are two classes (classes 1 and 2), six types, and several subtypes of CRISPR-Cas systems (1,7). The classification is primarily based on the Cas protein (class 2) or proteins (class 1) present within the effector module. Type II CRISPR typically encodes three Cas proteins, where Cas1 and Cas2 are required for spacer acquisition, and Cas9 acts as the entire effector module targeting and cleaving the foreign DNA (8). As antimicrobial employment of CRISPR only requires the recognition, binding, and cutting of the target, the most straightforward CRISPR-based tool consists only of Cas9 endonuclease and an editable CRISPR spacer array with tracrRNA (9). Once the complex scans the target locus and recognizes a protospacer adjacent motif (PAM) sequence, the nuclease activity of Cas9 mediates a double-stranded break in the target DNA that may result in cell death (10–13). The delivery of CRISPR antimicrobials into bacterial cells can be mediated by phages (phagemids) (10,14,15), or they can be introduced via utilizing natural bacterial strategies of horizontal gene transfer (HGT), such as conjugative plasmids (16–19). We have developed a CRISPR-based antimicrobial tool referred to here as CRICON (CRISPR via conjugation) (20). CRICON consists of two plasmids: a broad host range delivery plasmid RP4 and a mobilizable CRISPR plasmid (containing genetic sequences for CRISPR-Cas9 and tracrRNA) which transfers along with the delivery plasmid and mediates the nicking of the target sequence. CRICON was shown to efficiently reduce the prevalence of resistance in the *Escherichia coli* population harboring plasmid-carried *bla*CTX-M-14 and *bla*TEM-52b genes (20).

While antimicrobial tools employing CRISPR have demonstrated their proof-of-principle potential for AMR removal (16–18,21–23), they are usually applied in comparably short timescales. In addition, very little is known about the ecological and evolutionary factors that modulate their efficacy in more complex and natural environments. Indeed, natural microbial communities have a higher diversity of bacteria wherein mobile genetic elements, i.e., phages and plasmids, compete for suitable host cells. The benefit of plasmid uptake is highly dependent on the advantageous traits it can provide its host, e.g., antibiotic resistance. Although, the actual fitness effect of these opportunistic traits is determined by the surrounding environment. For example, resistance-conferring plasmids are more likely to be maintained under antibiotic exposure, while increased plasmid curing occurs in the absence of antibiotics (24). However, bacteria may exploit their defense or other mechanisms to resist unwanted genetic invasions. Since plasmid existence is essentially linked to host availability, plasmids readily adapt to changing host environments (25,26). Collectively, the ecological interactions responsively drive the evolutionary interplay between bacteria and their plasmids. Similarly, plasmid-host interactions are expected to affect the efficiency of conjugatively delivered antimicrobial tools, but for now, the longer-term evolutionary dynamics have not been studied.

Here, we investigated the evolutionary consequences of ESBL-harboring bacteria targeted by a CRISPR-based conjugatively delivered antimicrobial tool (CRICON) in a 10-day multispecies coevolution experiment. We set up synthetic microbial microcosms that simulate the gut microbiome with four ESBL-producing clinical *Escherichia coli* strains as CRICON targets (two of which harbor chromosomal and two with plasmid-based ESBL genes). We show that CRICON reduces the ESBL prevalence within the bacterial community, while the survival of ESBL-bacteria is highly dependent on the initial community composition. Interestingly, we observed that in one of the systems, the plasmid-based ESBL is maintained in the population by escaping into a more competitive dominating host species, *Klebsiella* sp. Further, we collected evolved ESBL-carrying *Klebsiella* sp. transconjugants and showed the increased conjugation rate of the ESBL plasmid and a partial resistance against the CRISPR-antimicrobials developed during the coevolution experiment. These results underline the importance of understanding both the ecological and evolutionary drivers within multispecies bacterial communities, which may disrupt the effective use of CRISPR-based antimicrobial strategies and, more generally, which may lead to unexpected and undesired outcomes of targeted therapies against multi-resistant bacteria.

## Materials and methods

### CRICON delivery strains and culture conditions

The CRICON plasmid system described in our previous study consists of two plasmids, a CRISPR plasmid and a delivery plasmid RP4 (20). The CRISPR plasmid (pCRISPR) was previously constructed from plasmid pCas9 (a gift from Luciano Marraffini, Addgene plasmid # 42876), encoding chloramphenicol resistance gene for selection and a CRISPR-Cas system including Cas9 endonuclease and a programmable CRISPR array site, mobilized via cloning of the *oriT* site (20). The ESBL-targeting CRISPR plasmid was programmed to target the *bla*CTX-M-15 gene by adding a specific CRISPR spacer at the CRISPR array site, as described earlier (20). This spacer-containing pCRISPR plasmid (pCas9-CTX-M-15_802) is referred to as S, while the CRICON control plasmid, without a CRISPR spacer (pCas9-oriT_ctrl), is referred to as CC. The CRISPR plasmids were transformed separately into *Escherichia coli* HB101 host strain carrying conjugative broad host range plasmid RP4 (carrying kanamycin and tetracycline resistance genes but with beta-lactamase gene *bla*TEM-2 knocked out; RP4-Λι*bla*TEM-2Δ^172−714^), hereafter referred to as RP4-Λι*bla*TEM (20) (**Figure 1a and Table 2**). The resulting strains carrying the plasmids were *E. coli* HB101 (RP4-Λι*bla*TEM) (pCas9-oriT), hereafter referred to as HB-CC (for **<u>C</u>**RICON **<u>c</u>**ontrol), and *E. coli* HB101 (RP4-Λι*bla*TEM) (pCas9-CTX-M-15_802), hereafter referred to as HB-S (for CRISPR **<u>s</u>**pacer targeting *bla*CTX-M-15 gene). Throughout this study, the bacteria were grown in 5 ml of Luria-Bertani (LB) broth (27) overnight at 37°C, 120 rpm. CRICON bacterial cultures were supplemented with 25 µg/ml kanamycin and 25 µg/ml chloramphenicol to maintain the carriage of both plasmids.

**Figure 1.**
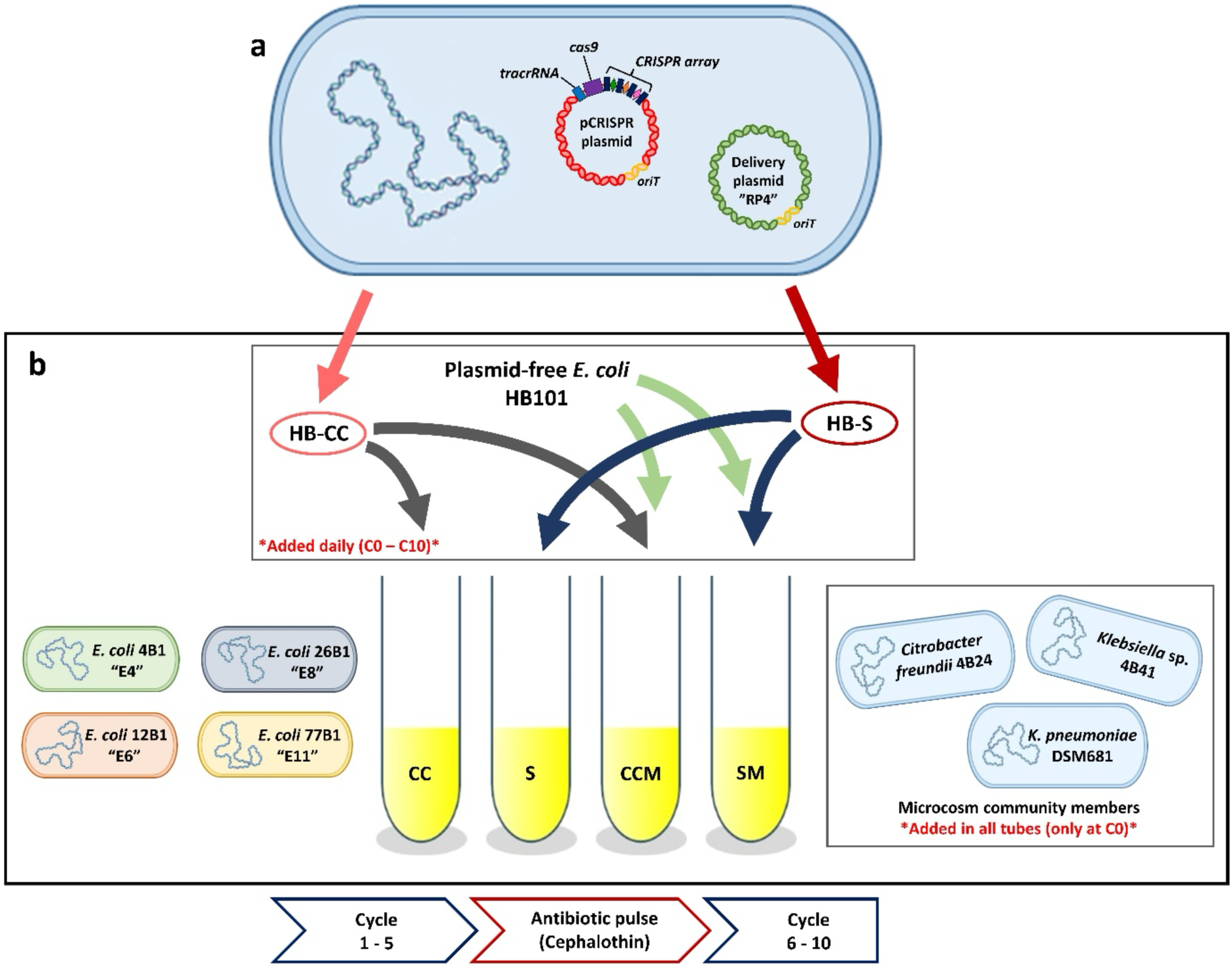
Bacterial setup and outline of the coevolution experiment. (a) the CRICON delivery strains used in this study. A control strain (HB-CC) harbors a mobilized CRISPR plasmid without a spacer, while a spacer-containing strain (HB-S) harbors a mobilized CRISPR plasmid with a spacer targeting *bla*CTX-M-15 gene. Both strains harbor RP4-Λι*bla*TEM plasmid as delivery plasmid mediating the conjugation. (b) The experimental design of the microcosm setups. Four CRICON-targeted *E. coli*-ESBL strains (E4, E6, E8, and E11) were tested individually in four different treatments (CC = CRICON control – without migration, S = CRICON spacer – without migration, CCM = CRICON control – with migration, and SM = CRICON spacer – with migration). CRICON strain was added daily into the communities. In the migration (M) treatments, plasmid-free *E. coli* HB101 was added daily. The microcosms also contained three other bacterial strains (*Citrobacter freundii* 4B24, *Klebsiella* sp. 4B41, and *K. pneumoniae* DSM681) as a core community. The cultures were refreshed daily (1:100) for 10 cycles (N = 3 replicates/treatment) with an antibiotic pulse of cephalothin after cycle 5 to select for the ESBL gene.

### Conjugation efficiency of the CRICON system into *E. coli*

To assess the conjugation efficiency of both control (CC) and spacer-containing (S) CRICON systems, overnight-grown CRICON donor (HB-CC or HB-S) and *E. coli* HMS174 (grown with 150 μg/ml rifampicin) recipient strains were diluted 1:100 in fresh LB broth and cocultured for 2 hours (4 replicates/coculture). Serial dilutions of the cocultures were plated on LB agar supplemented with 25 μg/ml kanamycin, 25 μg/ml chloramphenicol, and 150 μg/ml rifampicin to select only HMS174 transconjugants. Colony forming units (CFU) were determined from overnight plates grown at 37°C. The overnight-grown donor strains were also plated on LB agar supplemented with 25 μg/ml kanamycin and 25 μg/ml chloramphenicol to determine their cell number (CFU). The mean conjugation frequency was given as the ratio of formed transconjugants per initial donor cell (transconjugant CFU/ml: initial donor cell density).

### Pairwise conjugation test of CRICON to ESBL *E. coli* strains

The CRICON strains were initially tested for the efficiency in eradicating the *bla*CTX-M-15 gene in various *E. coli* ESBL strains isolated earlier from human fecal samples at Turku University Hospital (research permit T300/2020-1 of Hospital District of Southwest Finland). The conjugation was done as a coculture in 5 ml LB broth by adding 5 µl overnight-grown donor CRICON strain (either HB-CC or HB-S) and 5 μl of the 1:100 diluted overnight culture of the recipient ESBL strain (E4-E12) to obtain the initial ratio of 100:1 (donor: recipient) (**Table 1**). The cocultures were grown overnight at 37°C, 120 rpm in triplicates. The density of formed CRICON-ESBL transconjugants was determined by serial plating on LB agar supplemented with 150 μg/ml rifampicin, 50 µg/ml cephalothin, and 25 µg/ml chloramphenicol. CRICON efficiency was calculated as the final concentration of CRICON-ESBL transconjugants relative to the initial number of CRICON donor cells and compared between the CC and S treatments.

**Table 1.**
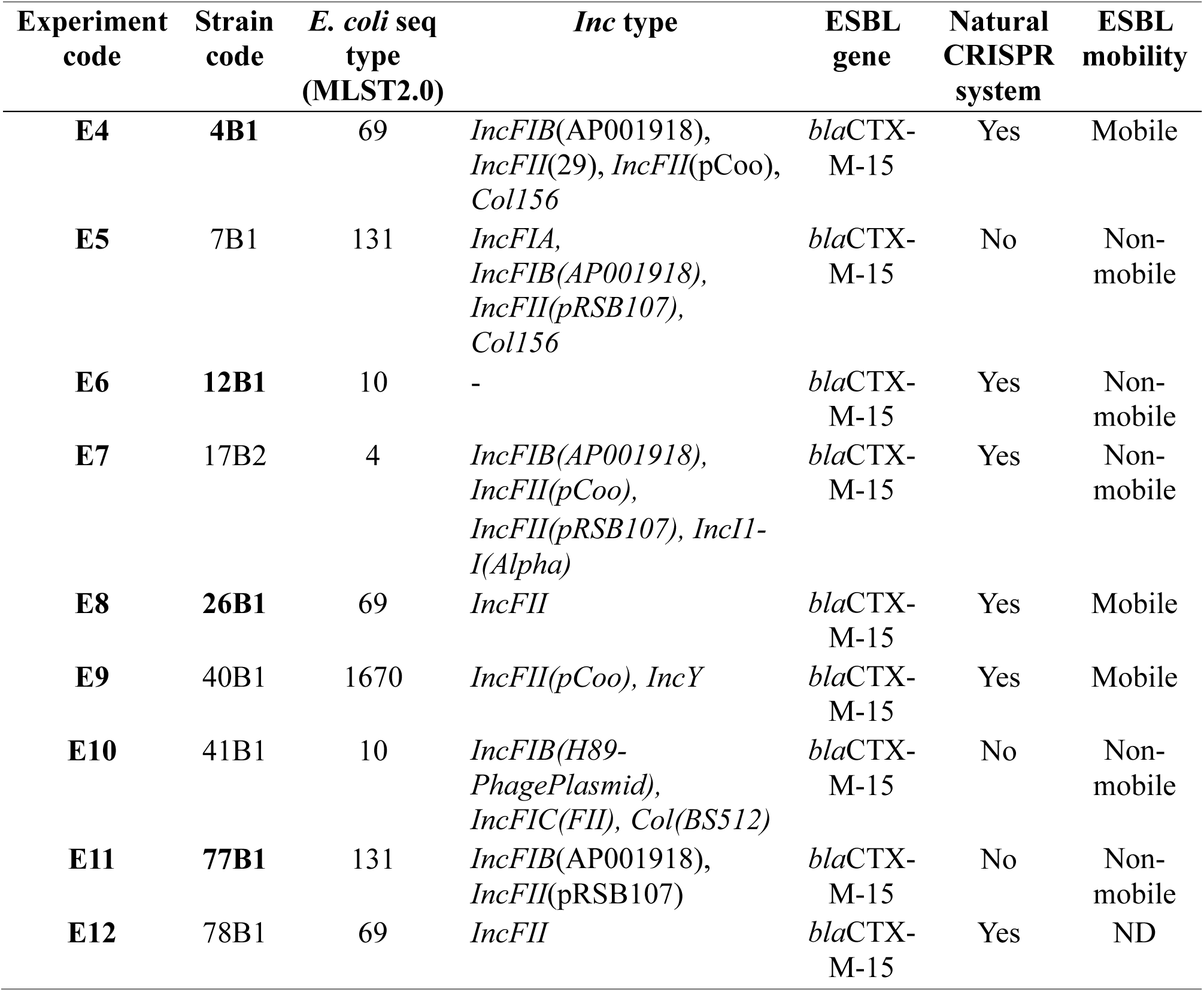
ESBL-*E. coli* strains used in this study as CRICON targets. ND = Not Determined.

**Table 2.**
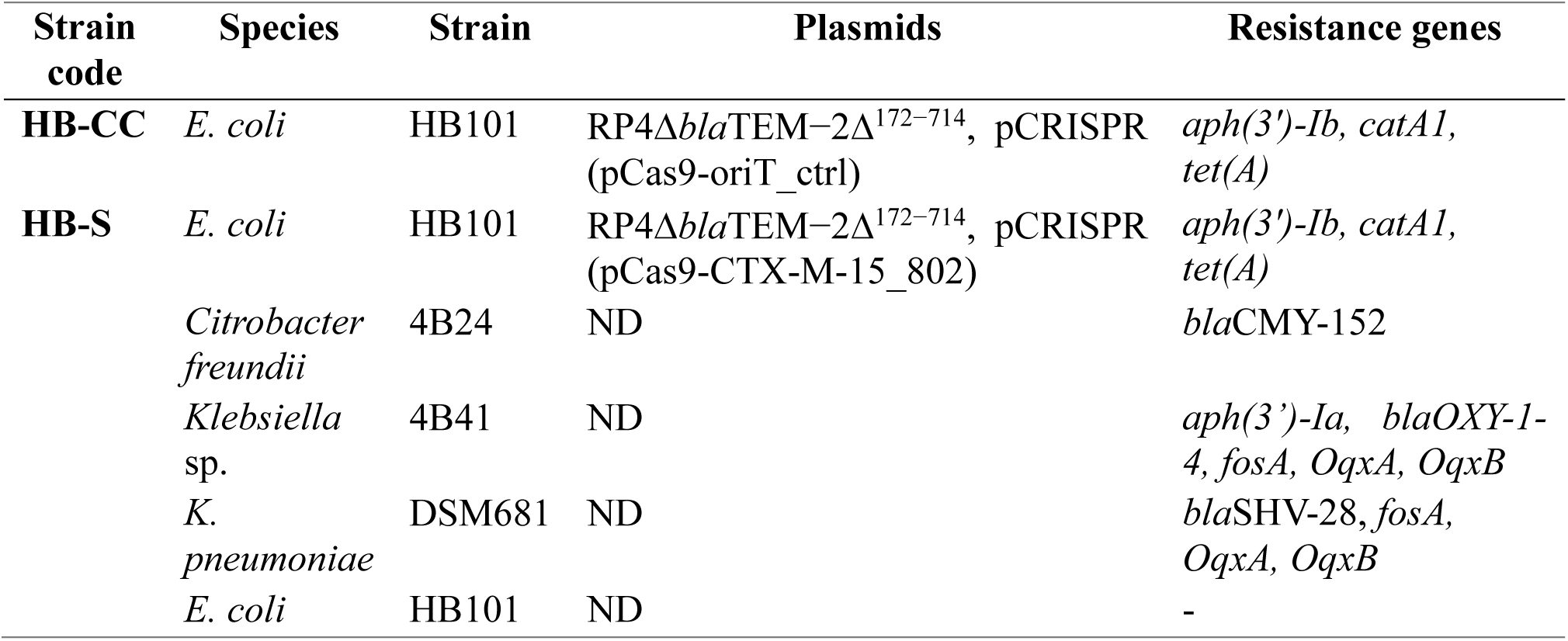
CRICON delivery strains and core community members used in the experiment. ND = Not Determined.

### Construction of synthetic bacterial community

Next, we assembled a synthetic bacterial community to test the efficiency of CRICON to remove ESBL within an ecologically more complex microbial environment. The core community included *Citrobacter freundii* 4B24, *Klebsiella* sp. 4B41 (both isolated from human fecal samples as referred above), and *K. pneumoniae* strain DSM681 (**Table 2**). Plasmid-free *E. coli* HB101 was used to assess if the migration of suitable donor cells affects the spread of CRICON and ESBL plasmids via conjugation. The community strains were grown in LB broth without antibiotics at 200 rpm, at 37°C overnight.

### Cross-inhibition test of community strains

All the community strains were tested for cross-inhibition against each other. 100 µl of the overnight culture of each strain was separately mixed with 3 ml of LB soft-agar and overlayed on LB agar plates without antibiotics (1 strain per plate). One milliliter of the remaining overnight culture was centrifuged at 5000×g for 1 min, and 10 µl of the supernatant was spotted on the bacterial lawn. The plates were incubated at 37°C overnight, and possible cross-inhibition was recorded as positive, faint, or no bacterial growth at the spotted area.

### Microcosm evolution experiment

The microcosm experiment was set up with four different settings (**Table 3**) based on two CRISPR systems (CC and S) and whether migrating plasmid-free cells were introduced into the community (M) during the experiment.

**Table 3.** Microcosm experimental settings.

| Treatment | Plasmid | Migration |
| --- | --- | --- |
| CC | CRICON control plasmid | Without plasmid-free cell migration |
| S | CRICON plasmid with spacer targeting *bla*CTX-M-15 gene | Without plasmid-free cell migration |
| CCM | CRICON control plasmid | With plasmid-free cell migration (*E. coli* HB101) |
| SM | CRICON plasmid with spacer targeting *bla*CTX-M-15 gene | With plasmid-free cell migration (*E. coli* HB101) |

**Note:** CC = control, S = spacer, M = migration

Four clinical ESBL-*E. coli* strains: 4B1 (E4), 12B1 (E6), 26B1 (E8), and 77B1 (E11) were chosen as CRICON targets in the microcosm experiment based on the preliminary test of CRICON efficiency. These strains possess different predicted plasmids and genetic associations of the ESBL gene (either plasmid-based or chromosomal) (**Table 1**). The ESBL target strains were cultivated in LB broth supplemented with 50 µg/ml cephalothin at 37°C, 200 rpm, overnight prior to the start of the experiment. Hence, the synthetic bacterial community consisted of five (no migration) or six (with migration) bacterial strains: one of the CRICON delivery strains (HB-CC control strain or HB-S targeting *bla*CTX-M-15 gene), one ESBL-*E. coli* strain (carrying *bla*CTX-M-15), the three core community members (*Citrobacter freundii* 4B24*, Klebsiella* sp. 4B41, and *K. pneumoniae* DSM681) and in the case of migration (M) treatments, the plasmid-free *E. coli* HB101. The phenotypic antibiotic resistance of the community strains was tested by cultivating bacterial culture overnight on cephalothin (50 µg/ml) and cefpodoxime-containing ESBL ChromID plates (Biomerieux) to confirm their specificity to allow only the growth of ESBL-*E. coli* strains.

Prior to the start of the microcosm experiment (cycle 0 or C0), 5 µl of overnight cultures of each community and ESBL strains were added to a 5 ml LB broth. 50 µl of CRISPR delivery strains (HB-CC or HB-S) and 5 µl of *E. coli* HB101 overnight-grown bacteria were added according to the treatments (**Table 1 and Figure 1b**). In total, 16 different microcosms (4 target ESBL strains in 4 treatments) were grown in triplicates for 16 hours at 37°C with 200 rpm agitation.

The next day, cycle 1 (C1) was started. Each culture was refreshed into fresh 5 ml LB broth by transferring 50 µl of the culture from the previous cycle. Then, 50 µl of HB-CC or HB-S, and 5 µl of HB101 (from fresh overnight cultures) were added according to the treatment (**Table 1 and Figure 1b**) and cultivated at 37°C, 200 rpm for 16 hours. The same procedure was repeated in the following days and the cultures were serially propagated for 10 cycles. An antibiotic pulse was applied between cycles 5 and 6 by adding cephalothin (to the final concentration of 50 µg/ml) to each microcosm (**Figure 3**).

### Sample collection and DNA isolation

One milliliter of each bacterial culture was harvested (after growing 16h, prior to the refreshing) from 6 timepoints: C1, C3, C5, C6, C8, and C10. The samples were stored at −80°C for DNA isolation. Additionally, another batch of the samples (1 ml/culture) was collected into a 2 ml cryotube containing 300 µl of 87% sterile glycerol to later isolate the evolved bacteria. The total DNA of all the samples was extracted with DNeasy® Blood & Tissue kit (Qiagen) following the manufacturer’s instruction for the QIAcube HT platform. The concentration of the samples was determined by Qubit 3.0 fluorometer using the dsDNA High Sensitivity kit (Invitrogen, ThermoFisher Scientific).

### Quantitative PCR (qPCR) with primers targeting *bla*CTX-M-15 gene

qPCR with probe assay was performed to follow the presence and abundance of the ESBL gene *bla*CTX-M-15. First, the qPCR probe (**Supplementary Table 1**) was designed to complement the *bla*CTX-M-15 gene in the samples. *bla*CTX-M-15-targeting primers (**Supplementary Table 1**) were optimized for efficiency before the experiment, and the crosscheck test against each community strain was done to ensure the specificity of the primers and probe to the target.

The qPCR final reaction volume of 20 µl contained 10 µl 2X SsoAdvanced Universal Probes Supermix (Bio-rad), 1 µl of 10 mM forward and reverse primers each, 0.5 µl of 10 mM probe, and 2 µl of DNA template (1 ng/µl) (**Supplementary Table 2**).

The 16S rRNA gene was used as a reference for normalization to calculate the relative abundance of the *bla*CTX-M-15 gene in the communities. qPCR for the 16S gene (**Supplementary Table 1 and 3**) was done without probe in a similar manner to the previous study (28).

All qPCR assays were performed with Bio-Rad CFX96 Touch™ Real-Time PCR Detection System (Bio-Rad), and the gene analysis was done with CFX Maestro™ 1.1 software (version 4.1.2433.1219, Bio-Rad). The relative abundance of *bla*CTX-M-15 gene from C1, C3, C5, C6 (for all microcosms), and C10 (for E8 and E11) was calculated by normalizing against 16S rRNA in the same samples and visualized with Rstudio (R version 4.3.0).

### Whole genome analysis of the community strains

Whole genome analysis of the community strains was also performed to check their defense systems, and plasmid assembly was conducted to map the mutations of the communities’ sequences after the evolution experiment. DNA was isolated using the Blood and Tissue DNA Isolation Kit (Qiagen) and Wizard® Genomic DNA Purification Kit (Promega). The strains were sequenced using the Illumina sequencing platform to produce paired-end sequencing reads of 150bp in length. Raw reads were trimmed with fastp (29) and assembled with spades (using option --isolate) (30). Acquired antibiotic resistance genes were identified from assembled contigs with ResFinder 4.5.0 (with minimum coverage of 60% and similarity threshold of 80%) (31–33) and possible plasmid carriage determined with PlasmidFinder (https://cge.food.dtu.dk/services/PlasmidFinder/) by using options Enterobacteriales.

### Metagenomic sequencing of the communities

Based on the qPCR result, the DNA of E8 and E11 samples cycles 1, 5, and 10 from migration treatment (CCM and SM) were proceeded to shallow shotgun metagenomic sequencing with DNBSEQ platform (PE150). Raw reads were trimmed with fastp (29) and the community structure determined with kraken v. 2.1.2 (34) using the standard database. Taxonomic groups provided by kraken output (*Klebsiella/Raoultella, Escherichia,* and *Citrobacter*) were used to produce a genus level estimate on the community composition, where *Klebsiella* group includes both *K. pneumoniae* and *K. sp.* strains, and *Escherichia* contains all ESBL, CRICON, and migration strains.

### Pairwise conjugation efficiency of E8 ESBL plasmid into community

The conjugation was initiated through overnight growth of donor and recipient strains in 5 mL LB broth supplemented with appropriate antibiotics (community strains: 150 μg/ml rifampicin, *E. coli* HB101: 25 μg/ml streptomycin, *E. coli* E8: 50 µg/ml cephalothin). Cocultures were started with the overnight culture of the donor and recipient transferred at 1:100 dilution into fresh LB broth and grown for 16 hours (4 replicates/culture). Dilutions of the cocultures were plated on LB agar supplemented with appropriate antibiotics (50 µg/ml cephalothin and 150 μg/ml rifampicin), selecting only for transconjugants. The density of formed transconjugants was determined as CFUs. The donor strains were plated on LB agar supplemented with 50 µg/ml cephalothin to determine their initial cell density. The mean conjugation frequency was calculated as the transconjugant cell density divided by the initial donor cell density.

### ESBL plasmid-carrier species determination from E8 community

Samples from the last cycle (C10) of E8 microcosms with migration treatment (SM and CCM) were spread onto ESBL plates (ChromID ESBL plate, Biomerieux) (100 µl, dilution 10^-3^) to isolate evolved ESBL-harboring strains. Eight single green colonies were picked per replicate per sample and re-streaked onto fresh ESBL plates. The re-streak was done twice to obtain pure isolates. These strains were then grown in 5 ml LB broth at 37°C, overnight without shaking. DNA was isolated from each strain (1 ml) with the Blood & Tissue kit (Qiagen) and used for fusion PCR that amplifies a fusion PCR product containing 16S rRNA (338F-534R) region and part of *bla*CTX-M-15 gene. The fusion PCR was used to confirm that each isolated strain was carrying the ESBL plasmid. Fusion-PCR reaction contains 25-30 ng of DNA, forward and reverse primers for *bla*CTX-M-15 at 10:1 ratio with the final concentration of 200 nm and the forward and reverse primers for 16S at 1:5 ratio with the final concentration of 250 nm (**Supplementary Table 1**) and 1x Phusion Master Mix with HF buffer in a total of 30 µl per one reaction. The amplification was done following the protocol mentioned in Dutra et al., (2022) and **Supplementary Table 4**. The genomic DNA from the strains positive for the *bla*CTX-M-15 gene were then prepared for partial 16S rRNA gene Sanger sequencing (ABI Prism® 3130 xl sequencer, Applied Biosystems Hitachi) for taxonomic identification with 338F primer.

### Growth rate analysis of ESBL-*Klebsiella* sp. transconjugants

Growth rate analysis was performed on the transconjugants identified as ESBL-*Klebsiella* sp. as well as the community strains to determine their growth parameters (growth and yield). Overnight cultures were initiated in LB broth (transconjugants and *E. coli* ESBL strains: 50 µg/ml cephalothin, community strains (*Citrobacter freundii* 4B24*, Klebsiella* sp. 4B41, and *K. pneumoniae* DSM681): 150 μg/ml rifampicin, CRICON strains: 25 μg/ml kanamycin and 25 μg/ml chloramphenicol, *E. coli* HB101: 25 μg/ml streptomycin) before dilution of 1:100 with fresh LB medium. The growth of the bacterial strains was determined with a Bioscreen C MBR machine (Bioscreen, Oy Growth Curves Ab Ltd.) at 37°C, low shaking. The absorbance was measured at OD_600_ in 5-min intervals for 24h. The growth curves, growth rate (R), and maximum yield (K) were calculated from the data using RStudio (R version 4.3.0), with R source code (see Jonsdottir et al., 2023), based on a previously described MATLAB code (36).

### Assessment of the transfer rate of original (pE8) and evolved E8 (pE8-*Klebsiella* sp.) ESBL plasmid into *E. coli*

To allow for the same genetic host background of the evolved E8 plasmids from *Klebsiella* sp. ESBL transconjugants (pE8-*Klebsiella* sp.) and the original E8 plasmid (pE8), they were first conjugated into the same host: *E. coli* HB101. The conjugation was initiated through overnight growth of ESBL donors and recipient strains in 5 mL LB broth supplemented with appropriate antibiotics (transconjugants and *E. coli* E8: 50 µg/ml cephalothin, *E. coli* HB101: 25 μg/ml streptomycin). Cocultures were started with the overnight culture of ESBL donor and recipient strains transferred at 1:100 dilution into fresh LB broth and grown for 4 hours. ESBL-transconjugants were selected by plating on LB agar supplemented with 50 µg/ml cephalothin and 25 μg/ml streptomycin. Once all plasmids were successfully conjugated into *E. coli* HB101, further conjugation was done into *E. coli* HMS174 to determine the plasmid transfer rate. The conjugation was initiated through overnight growth of the HB101 donor (harboring either the original (pE8) or evolved ESBL (pE8-*Klebsiella* sp.) plasmid) and recipient HMS174 strains in 5 mL LB broth supplemented with appropriate antibiotics (ESBL-transconjugants: 50 µg/ml cephalothin, *E. coli* HMS174: 150 μg/ml rifampicin). Cocultures were started with the overnight culture of the donor and recipient transferred at 1:100 dilution into fresh LB broth without antibiotics and grown for 4 hours (4 replicates/culture). Dilutions of the cocultures were plated on LB agar supplemented with 50 µg/ml cephalothin and 150 μg/ml rifampicin, selecting only for HMS174 ESBL-transconjugants. Colony forming units (CFU) were then determined. The donor strains were plated on LB agar supplemented with 50 µg/ml cephalothin to determine their cell number (CFU). The mean conjugation frequency was given as the transconjugant cell density divided by the initial donor cell density.

### Efficiency of CRICON system against the evolved plasmid from E8 ESBL**-***Klebsiella* sp. transconjugants (pE8-*Klebsiella* sp.)

After coculturing, the CRICON efficiency against the evolved ESBL carriers was assessed via pairwise conjugation tests and subsequent qPCR. The conjugation was initiated through overnight growth of donor and recipient strains in 5 mL LB broth supplemented with appropriate antibiotics (*E. coli* CRICON strains: 25 μg/ml kanamycin and 25 μg/ml chloramphenicol, transconjugants: 50 µg/ml cephalothin). Cocultures were started with the overnight culture of the donor and recipient transferred at 1:100 dilution into fresh LB broth and cultivated for 4 hours (4 replicates/culture). qPCR with probe assay quantifying the ESBL gene (*bla*CTX-M-15) (described above) was performed to determine the gene abundance with 16S as reference.

## Results

Carriage of ESBL-harboring bacteria in the gut can compromise the antibiotic treatment. This has prompted innovation in new approaches, such as utilizing CRISPR systems, to limit the spread and carriage of ESBL. We previously developed a simple two-plasmid system where conjugative plasmid RP4 delivers a CRISPR-plasmid to ESBL-carrying recipient bacteria (20). In pairwise conjugation experiments, this system, designated as CRICON, shows potential in reducing the prevalence of ESBL genes, but the experiments in more complicated setups are lacking. Therefore, we here set out to investigate the *in vitro* survival of ESBL-genes carried by *E. coli* in a multispecies synthetic community. In this community, a *Klebsiella* sp. strain tends to dominate the microbial system. Thus, we aimed to determine whether ESBL-genes can survive despite the survival of their original *E. coli* host. To ensure the maintenance of CRICON plasmids in the community, the system was continuously supplemented with CRICON-carrying bacteria.

### Two-plasmid conjugative anti-ESBL CRISPR-system (CRICON) successfully removes ESBL from several clinical *E. coli* strains

First, we investigated if the selected nine *E. coli* strains originating from human fecal samples could receive the CRICON plasmids and if the CRICON system functions efficiently, which was seen as the decreased abundance of ESBL genes. The transfer rates of CRICON CC and S were first confirmed to be similar (Supplementary Table 5). All the strains were previously determined to carry an ESBL gene *bla*CTX-M-15, and due to their human origin, they would thereby represent realistic targets for CRICON-induced eradication of ESBL.

CRICON system targeting *bla*CTX-M-15 gene (HB-S) decreased the number of ESBL transconjugants in several ESBL strains tested as compared to the CRICON control strain HB-CC (lacking a spacer) (**Supplementary Table 6**). CRICON was confirmed efficient against E4, E6, E8, E9, E11, and E12. However, in E5, E7, and E10, none or only a marginal number of formed transconjugants were observed in either of the CRICON treatments, indicating that some ESBL strains were not efficiently compatible with the CRICON plasmid uptake. Combining this result with the characteristics of each ESBL strain shown in **Table 1**, four strains were chosen for the evolution microcosm experiment to study how the ecological factors affect the efficiency of CRICON to operate in a multispecies environment (**Table 1**, highlighted in bold).

### Cross-inhibition between strains

To investigate whether the strains naturally suppress or kill the other strains in the microcosm experiment, we plated the supernatant of the bacterial cultures on the lawns of all other strains. E4 showed positive inhibition on HB-CC and unclear results on HB-S and HB101, whereas E8 showed positive inhibition against HB-S, HB-CC, and HB101 (**Supplementary Table 7**). Other strains showed no inhibition (**Supplementary Table 7**). We decided to include the inhibiting strains in the experimental setup to potentially cast some light on the role of pairwise inhibiting effects on CRICON efficacy.

### Abundance of *bla*CTX-M-15 throughout the multispecies microcosm experiment

We set up a 10-day evolutionary experiment consisting of a synthetic bacterial community and a single ESBL (*bla*CTX-M-15) positive *E. coli* strain. Further, the communities were supplemented with *E. coli* HB101 strain harboring a CRICON-system that either carries (S) or does not carry (CC) the *bla*CTX-M-15 targeting spacer. Two of the experimental setups were also continuously introduced with CRICON-free HB101 strain (M, i.e., migration) to simulate the presence of potential plasmid-free hosts for either the CRICON system or ESBL plasmids. We hypothesized that such hosts may help either or both plasmid systems prevail in the system. Also, we considered the possibility that the absence of antibiotics may naturally cure the system of ESBL-genes. As such, we decided to introduce a single pulse of antibiotic cephalothin to the microcosms after 5-days of serial culturing. The community setups and the evolutionary experiment are depicted in **Figure 1**.

Quantitative PCR (qPCR) was utilized to measure the abundance of the *bla*CTX-M-15 gene during the evolutionary experiment in the multispecies microcosms. After normalizing the results with 16S gene abundance, our results show that the *bla*CTX-M-15 gene disappeared in E4, E6, and E11 communities both from the control (CC) and spacer (S) treatment. This suggests that in these systems, the community dynamics drove the ESBL disappearance rather than the targeted CRICON treatment (**Figures 2** and **3**). However, in the E8 communities, the CRICON treatment decreased the relative abundance of the *bla*CTX-M-15 gene from the community compared to CC and S treatments (**Figures 2** and **3**). Neither the migration of plasmid-free cells nor the antibiotic pulse (at day 5) affected the *bla*CTX-M-15 survival.

**Figure 2.**
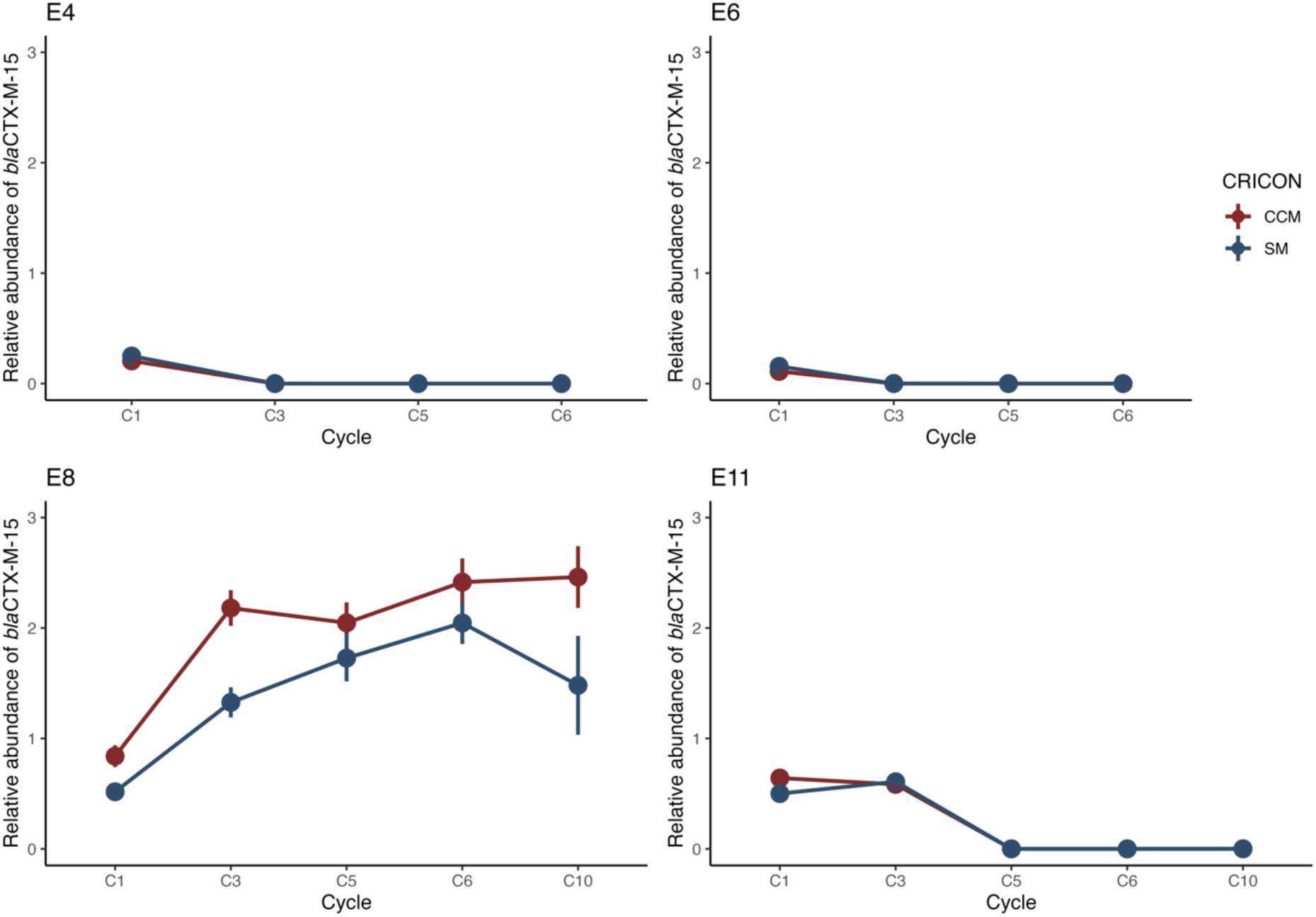
The relative abundance of *bla*CTX-M-15 during the 10-day evolution microcosm experiment in treatment with migration (M). E4 and E8 communities possess a mobile ESBL gene, and E6 and E11 a non-mobile ESBL gene. CCM = CRICON control with the migration of plasmid-free host cells, and SM = CRICON spacer treatment with the migration of plasmid-free host cells.

**Figure 3.**
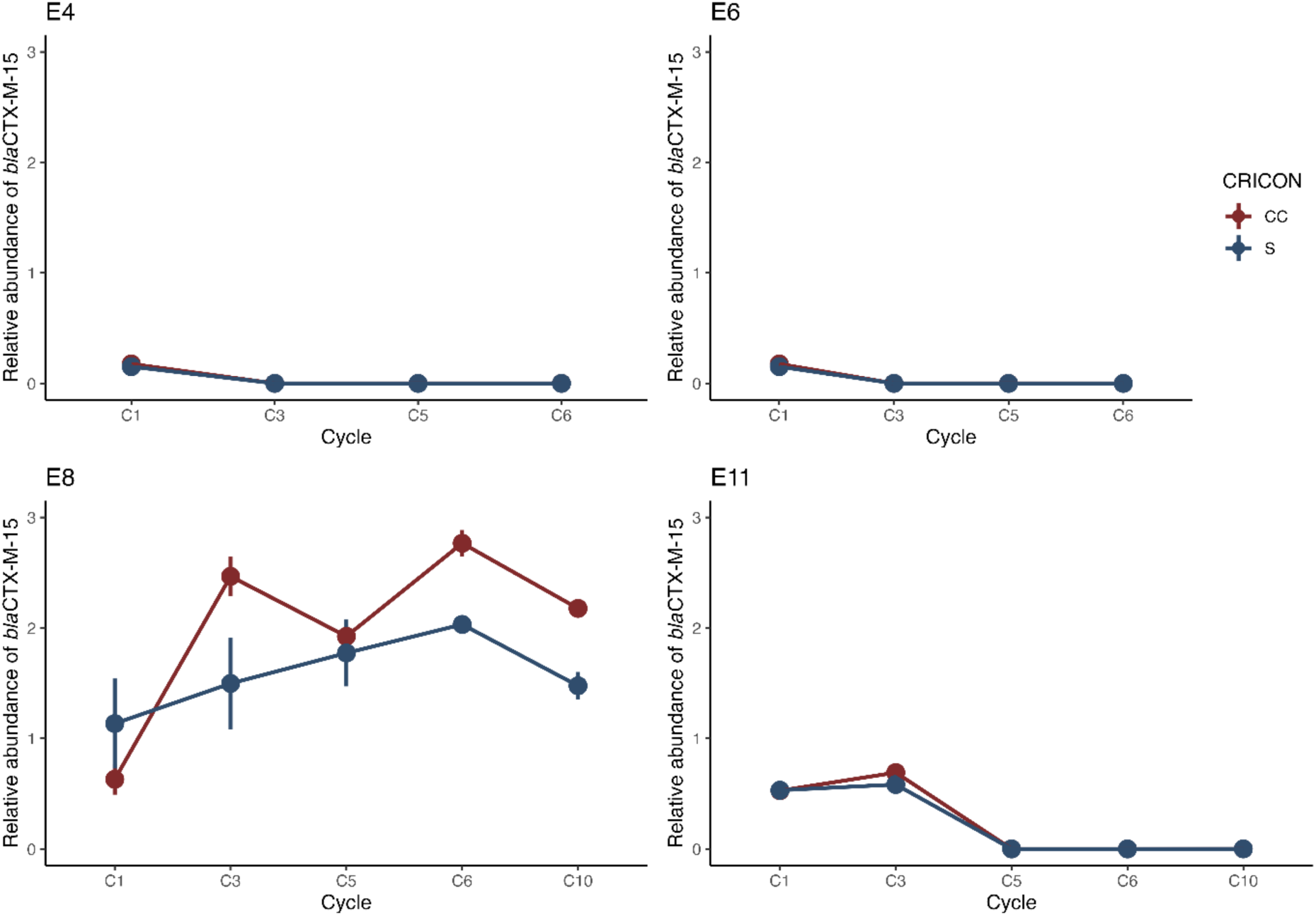
The relative abundance of *bla*CTX-M-15 during the 10-day evolution microcosm experiment without migration. E4 and E8 communities possess a mobile ESBL gene, and E6 and E11 a non-mobile ESBL gene. CC = CRICON control, S = CRICON spacer treatment, without the addition of plasmid-free host cells.

### Community composition throughout the microcosm experiment

The microcosm experiments showed that only the communities with E8 ESBL-strain could retain the *bla*CTX-M-15 throughout the experiment. To determine whether this was due to the dominance of the original E8 strain, we isolated twenty-four clones from the last cycle (C10) of migration treatments CCM and SM of the E8-containing experiment. Samples were plated on ESBL plates, and the resistant clones were picked up. The resistance tests of the original community strains showed that only the target ESBL-*E. coli* strains (E4, E6, E8, and E11) were able to grow on cefpodoxime-containing ESBL and cephalothin plates (except for strain 4B24, which grew on cephalothin). However, the bacterial lawns spread on chromogenic ESBL plates showed turquoise colonies (an indicator of *Klebsiella* genus) dominating all the E8 communities, indicating that *E. coli* (brown colonies on ESBL plates) was no longer the dominant ESBL carrier species within these communities. To exclude spontaneous resistance development via, e.g., mutations, all the clones were shown to carry *bla*CTX-M-15 via fusion-PCR. 16S rRNA sequencing of the clones revealed that all isolates were *Klebsiella* sp. 4B41, not the original *E. coli* E8 strain nor the *K. pneumoniae* strain involved in the community (**Supplementary Figure 1**). In other words, the survival of *bla*CTX-M-15 in E8 communities was most probably due to the transfer of ESBL gene-containing mobile genetic element to *Klebsiella* sp. 4B41.

To further determine the ecological dynamics within the experimental setups, we conducted metagenomic sequencing at different time points of the experiment (Supplementary Figure 2). Interestingly, *Klebsiella* was shown to be the most abundant genus in all the communities after cycle 1. It also remained the dominant strain throughout the experiment. These two results suggested that, indeed, *Klebsiella* took over the systems and only in E8-containing experiments, it became the leading player to harbor the ESBL-containing element.

### E8 conjugative transfer to potential recipients and subsequent growth

Given that the ESBL gene was consistently retained in communities with the E8 strain, we decided to investigate the conjugation dynamics of the E8 strain. E8 was cocultured with the available recipient for 16 hours, and the number of transconjugant cells was measured. Transconjugant *Klebsiella* sp. 4B41 cells were almost an order of magnitude more abundant than *E. coli* HB101 or two orders of magnitude more abundant than *K. pneumoniae* DSM681 cells after pairwise conjugation and subsequent growth (**Table 4**).

**Table 4.** Average transfer rate of E8 ESBL plasmid to synthetic community members.

| Recipient strain | CFU/mL | CFU/mL per donor cell |
| --- | --- | --- |
| *E. coli* HB101 | 1.445 x 10<sup>7</sup> | 1.225 x 10<sup>-2</sup> |
| *Klebsiella* sp. 4B41 | 1.26 x 10<sup>8</sup> | 1.068 x 10<sup>-1</sup> |
| *K. pneumoniae* DSM681 | 5.65 x 10<sup>6</sup> | 4.788 x 10<sup>-3</sup> |

### Growth analysis of E8 plasmid in *Klebsiella* sp. (E8 transconjugant clones)

The high number of ESBL-positive *Klebsiella* sp. cells after conjugation and overnight culturing suggest that the growth dynamics of *Klebsiella* sp. might be significantly higher than other strains in the experiment. This was true to some extent as *Klebsiella* sp. 4B41 clones isolated from CCM experiments produced the highest yield (**Table 5**). The growth rate of E8 transconjugant *Klebsiella* sp. 4B41 clones were similar between CCM and SM treatment (**Figure 4 and Table 5**). The growth rate and yield of other strains in the experimental setups were similar to *Klebsiella* sp. 4B41, besides the CRICON strains (HB-CC and HB-S) and plasmid-free HB101.

**Figure 4.**
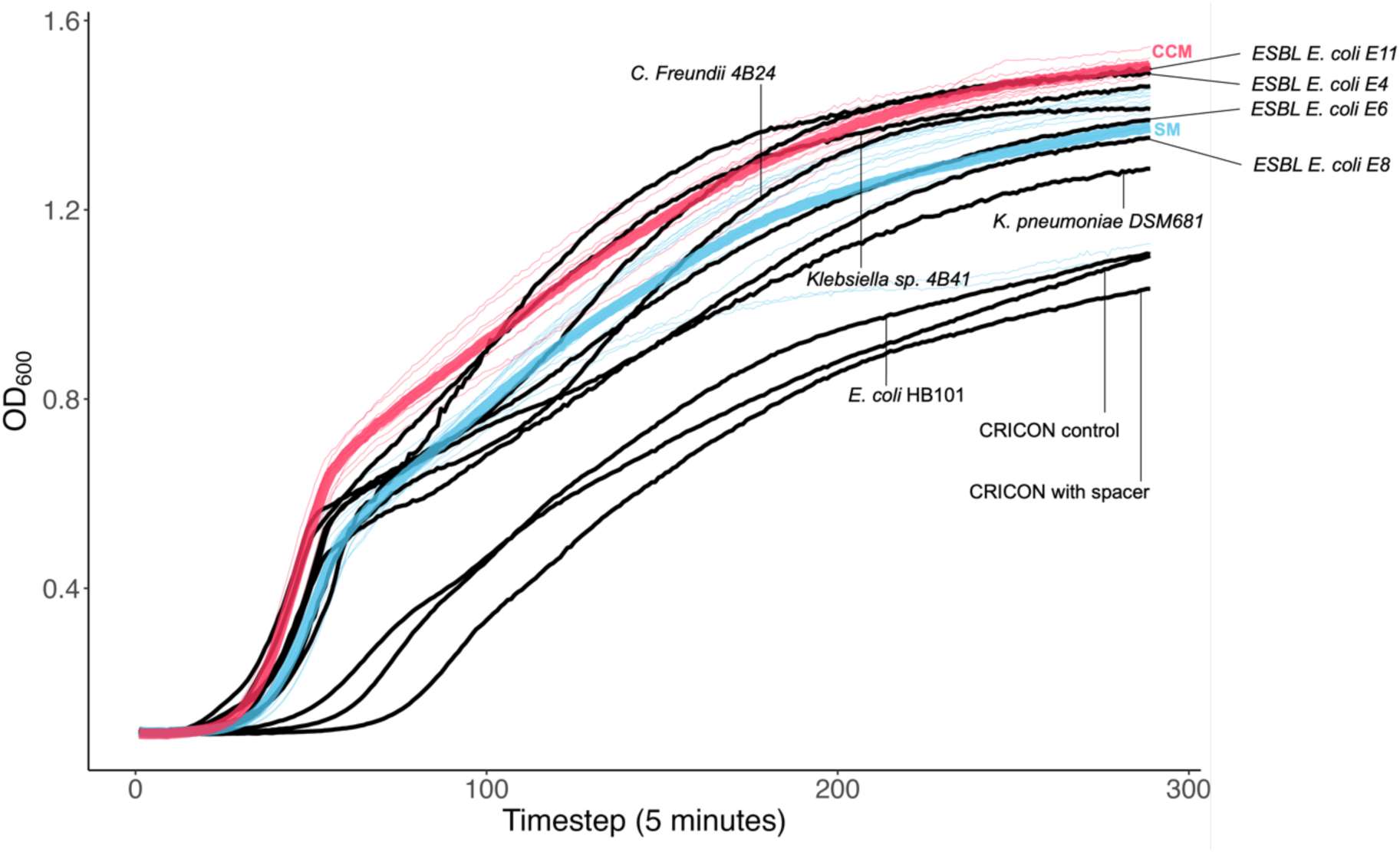
Growth curves of E8 ESBL clones from both CRICON control (CCM; red) and CRICON spacer (SM; blue) treatment at the microcosm endpoint (*Klebsiella* sp. (pE8) transconjugants). Additionally, growth curves of core community members (*C. freundii* 4B24, *Klebsiella* sp. 4B41, and *K. pneumoniae* DSM681), CRICON strains (control and spacer), targeted ESBL *E. coli* strains (E4, E6, E8, and E11) and *E. coli* HB101 (host of CRICON system) can be seen in black. Growth rate and maximum yield were calculated based on this data and can be seen in Table 5.

**Table 5.** Growth rate and maximum yield of the strains involved in this study and the microcosm endpoint *Klebsiella* sp. (pE8) transconjugants.

| Strain | Growth rate (r) | Maximum yield (K) |
| --- | --- | --- |
| *Klebsiella* sp. (pE8-CCM) clones | $0.014 \pm 0.0003$ | $1.49 \pm 0.0189$ |
| *Klebsiella* sp. (pE8-SM) clones | $0.012 \pm 0.0009$ | $1.36 \pm 0.1234$ |
| *C. freundii* 4B24 | 0.012 | 1.41 |
| *Klebsiella* sp. 4B41 | 0.012 | 1.45 |
| *K. pneumoniae* DSM 681 | 0.013 | 1.28 |
| *E. coli* HB101(plasmid-free) | 0.008 | 1.09 |
| *E. coli* HB101 (CC; CRICON control) | 0.007 | 1.08 |
| *E. coli* HB101 (S; CRICON with spacer) | 0.008 | 1.02 |
| ESBL *E. coli* E4 | 0.011 | 1.48 |
| ESBL *E. coli* E6 | 0.011 | 1.38 |
| ESBL *E. coli* E8 | 0.012 | 1.34 |
| ESBL *E. coli* E11 | 0.011 | 1.49 |

### Conjugation of original pE8 plasmid (pE8) vs. evolved pE8 (pE8-*Klebsiella* sp.) into *E. coli*

Given the successful transfer of ESBL-gene to *Klebsiella* sp. in E8-containing communities, we attempted to determine if the mobile element had indeed remained mobile and improved its ability to disseminate. Hence, we assessed the transfer rate of the original E8 ESBL plasmid (pE8) and the evolved plasmid carried by the *Klebsiella* sp. (pE8-*Klebsiella* sp.) clones isolated from the endpoint of the experiment (C10). In order to remove the evolved growth characteristics of the plasmid hosting *Klebsiella* sp. clones, the plasmids were first transferred to an intermediate host, *E. coli* HB101. The subsequent conjugation experiments were done between HB101 carrying pE8 or evolved pE8-*Klebsiella* sp. plasmids and another *E. coli* strain (HMS174). The results show that the conjugation rate into new recipient cells was significantly higher in the evolved pE8 plasmid (pE8-*Klebsiella* sp.), with a difference of two orders of magnitudes (CFU/mL), compared to the original pE8 plasmid. However, no difference was observed between pE8-*Klebsiella* sp. plasmids evolved in the CC or S treatments. To conclude, the increased plasmid transfer rate was more likely due to coevolution with the new *Klebsiella* sp. host rather than a response from being targeted with CRICON (**Figure 5**).

**Figure 5.**
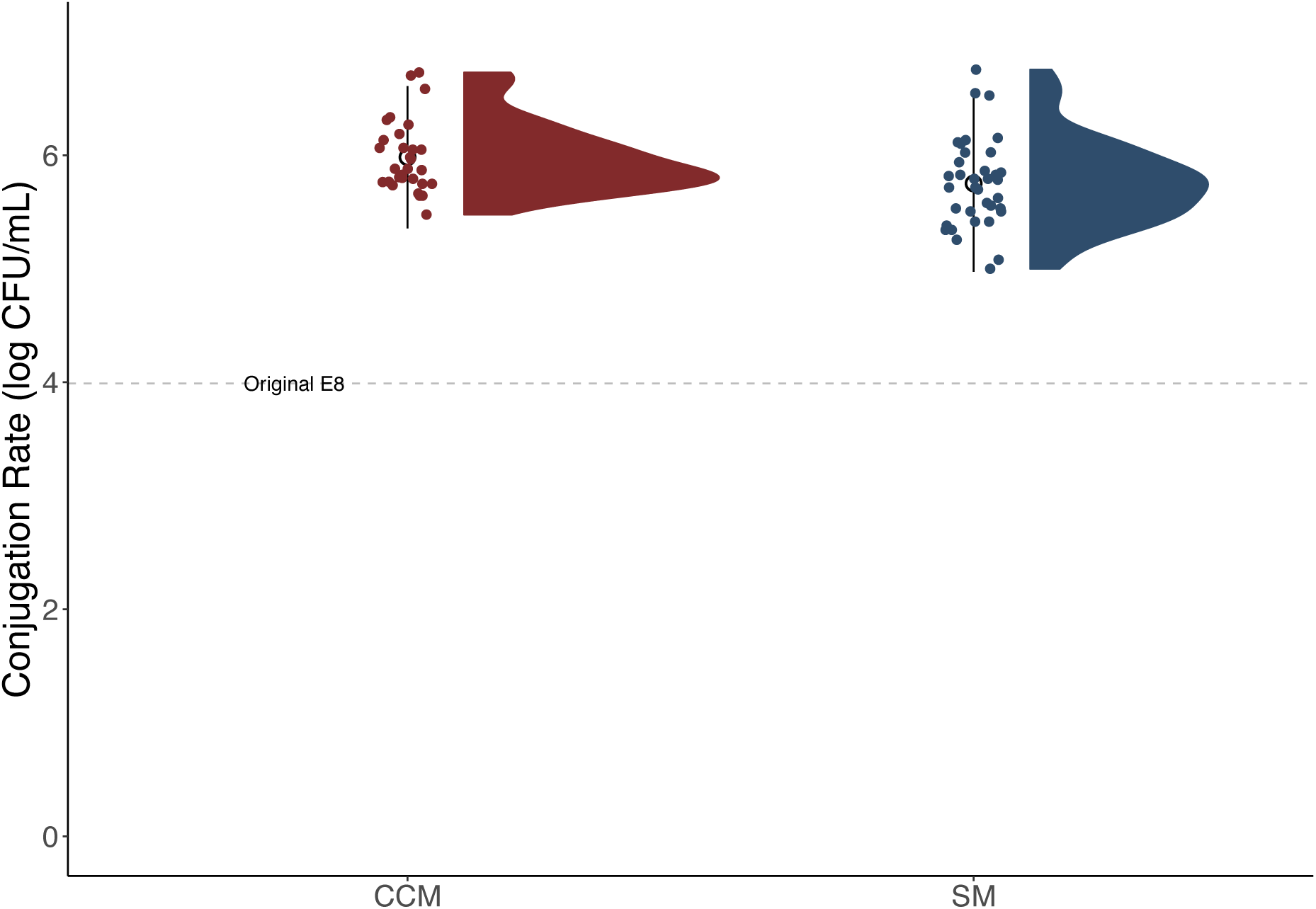
Conjugation rate of original and evolved pE8 ESBL plasmids. Original pE8 derived from *E. coli* E8 strain and evolved pE8 plasmids (pE8-*Klebsiella* sp.) from the *Klebsiella* sp. clones collected from the endpoint of the microcosm experiment. The evolved pE8-*Klebsiella* sp. conjugation rates are distinguished by treatment of CRICON control (CCM; red) and CRICON spacer (SM; blue) as well as the original E8 ESBL plasmid (pE8, grey dashed line). All the plasmids were conjugated from *E. coli* HB101 into *E. coli* HMS174 to be able to compare using the same genetic background.

### CRICON efficiency against the evolved *Klebsiella* sp. (pE8) transconjugant clones

Altogether, the presence of the CRICON system (as compared with or without ESBL gene targeting spacers) decreased the relative abundance of *bla*CTX-M-15, yet the effect seemed minor and not sufficient for efficient ESBL clearance. Hence, we wanted to examine if the presence of spacers had any impact on the ESBL prevalence now mainly carried by the new host *Klebsiella* sp. The efficiency of the CRICON system was assessed against the evolved *Klebsiella* sp. (pE8) clones isolated from the endpoint of E8 communities via a conjugation experiment between the clones and the CRICON delivery strains (HB-CC and HB-S). The CRICON efficiency was subsequently measured with qPCR analysis as *bla*CTX-M-15 relative abundance. The results showed that after a 4-hour conjugation with the CRICON delivery strains, 60% of *Klebsiella* sp. (pE8) transconjugant clones that evolved under the control treatment (CCM) (6 out of 10 clones) were still sensitive to the spacer treatment as less *bla*CTX-M-15 gene were detected (red dots, negative Δ values) than the spacer treatment (blue dots, negative Δ values) (**Figure 6**). On the other hand, most of the *Klebsiella* sp. (pE8) transconjugant clones that evolved under the SM treatment (9 out of 12) showed increased resistance towards the CRICON treatment as they maintained a higher abundance of the *bla*CTX-M-15 gene (blue dots, positive Δ values) compared to the control (red dots, positive Δ values) (**Figure 6**).

**Figure 6.**
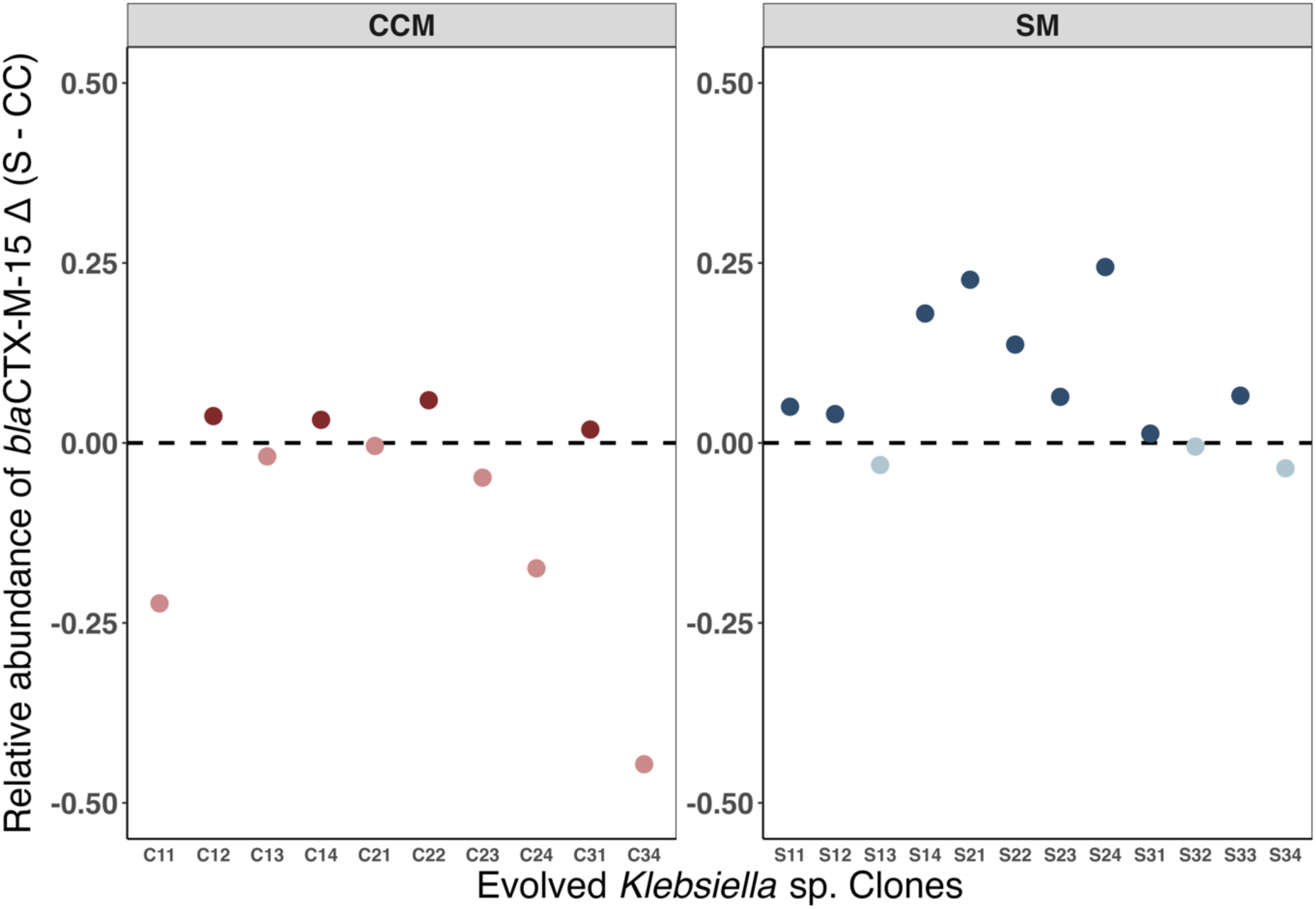
CRICON efficiency against *Klebsiella* sp. (pE8) transconjugants evolved under CRICON control (CCM) or CRICON spacer (SM) treatment. CRICON efficiency is determined as a difference in ESBL gene relative abundance (*bla*CTX-M-15^S^– *bla*CTX-M-15^CC^ = Δ*bla*CTX-M-15) in *Klebsiella* sp. (pE8) transconjugant clones. The plots are separated by the CRICON treatment during the microcosm experiment, CRICON control (CCM; red dots, left), and CRICON spacer (SM; blue dots, right).

## Discussion

The presence of antibiotic-resistant bacteria in the gut poses serious healthcare challenges. One of the main contributors to the problem is ESBL-*E. coli* strains that are resistant to a wide range of commonly used beta-lactam antibiotics. In this study, we investigated the survival of gut-derived ESBL-*E. coli* in a synthetic multispecies community that was continuously supplemented with *E. coli* strain (HB101) that carried CRICON, targeting the ESBL-gene (*bla*CTX-M-15). In our setup, the communities became heavily dominated by *Klebsiella* sp. strain, which could have caused the remarkable reduction of ESBL-*E. coli* in most of the systems (E4, E6, and E11). Sünderhauf et al. (2023) showed that community competitors affect the cost of plasmid carriage and maintenance, and that adding growth partners can drive plasmid loss from focal species in a short time frame. The competition pressure from other members of the communities, in this case, *Klebsiella* sp. 4B41 may drove the loss of the ESBL-*E. coli* rather than the CRICON treatment. However, in E8 communities, the ESBL gene persisted in the system although CRICON treatment reduced the prevalence of *bla*CTX-M-15 to some extent. Our results demonstrate that the interaction between the community members can essentially affect the persistence of ESBL plasmid in the system and the ability of CRICON to remove the ESBL gene efficiently. The result from the cross-inhibition test further supports this possibility, as only E8 showed inhibition against the CRICON strains HB-S, HB-CC, and migration strain HB101. The host strain and the mobility of its *bla*CTX-M-15 gene might also play a role in how the *bla*CTX-M-15 gene persisted in E8 communities since all ESBL-*E. coli* strains used in the experiment with the non-mobile ESBL gene disappeared from the system as early as cycle 2. However, the migration of plasmid-free cells in the systems did not affect the ability of CRICON to remove the *bla*CTX-M-15 gene from the system.

Further investigation into E8 communities by looking at the clones isolated from the endpoint microcosm showed that mobile ESBL-plasmid was transferred into the dominating *Klebsiella* sp. strain, thus allowing the system to retain the antibiotic resistance despite the rapid decrease of its original *E. coli* host. E8 ESBL strain showed the highest conjugation frequency to *Klebsiella* sp. compared to other strains in the system, providing the ESBL plasmid a pathway to escape to more competitive hosts in order to remain in the system. Further, the conjugation frequency of the plasmid in the E8 strain (pE8) was also higher when compared to the other mobile ESBL-carrying strain E4 (data not shown). Interestingly, the conjugation rate of the evolved pE8 ESBL plasmids carried by *Klebsiella* sp. transconjugant clones (pE8-CCM and pE8-SM) developed even higher than the original pE8 plasmid. These results show that the pE8 ESBL plasmid was initially more compatible with the synthetic community regarding its host range and transfer capacity. This, in turn, may have enabled its evasion of the CRICON treatment and efficient persistence within the microbial community.

The evolution of pE8 in different CRICON treatments resulted in another interesting outcome; while the *Klebsiella* sp. (pE8-CCM) clones (evolved without CRICON targeting) were still prone to CRICON, the *Klebsiella* sp. (pE8-SM) transconjugant clones exposed to the CRICON spacer treatment showed decreased CRICON efficiency in almost all clones. The escape from CRISPR-antimicrobial tools may appear via point mutations occurring under the positive selection for the AMR gene (12,23,38), via anti-CRISPR gene selection (39,40), rearrangements of the conjugative plasmid sequences (17), or the mutations in *cas* genes or spacer arrays, resulting in inactivation of CRISPR-Cas system and resistance evolution (20,22,41). Even though our study did not explore the genetic changes in the evolved plasmids, these results support the idea that certain level of resistance against CRISPR antimicrobials may develop.

Many factors can affect the efficiency and outcome of CRISPR tools for removing the ESBL gene, especially in a complex community with diverse microbes, for example, the conjugation efficiency, fitness costs, barriers to plasmid uptake and establishment, the mobility and copy number of targeted gene/plasmid, the resistance of targeted genes (e.g., through mutation, inhibition by toxin-antitoxin, anti-CRISPR), type of CRISPR-Cas system to targeted strains, multiplexing (i.e., targeting multiple sequences simultaneously), and the selection of delivery vectors (6,42). Suitable donors should also be considered for the target community, as well as the cost of CRICON plasmid and the expression and cytotoxicity of Cas9 in different species. Broad host range plasmids are a potential option for conjugative delivery of genetic tools (43) also due to their large coding capacities (44). Additionally, no recipient factors for DNA uptake have been identified (45), which in principle means, that recipient cells cannot avoid the uptake of the DNA via conjugation. Moreover, the transconjugants can also become donors for the subsequent re-conjugation (17,46). However, the efficient transfer rate and plasmid maintenance of any conjugative CRISPR tool are essential for successful removal of target genes from the microbial population. Further, these factors also determine the survival of the targeted gene within the community when located in a native conjugative (ESBL) plasmid. When looking at the efficiency of the CRICON system in our study, both control (HB-CC) and spacer (HB-S) strains exhibit similar conjugation frequency, yielding similar transfer efficiency of both strains into new recipient cells. However, they both showed the lowest growth rate and yield compared to other strains in the communities, and both lost their activity after a short period (data not shown). The loss of CRISPR activity was also shown in various other studies, even under strong selective pressure of antibiotic selection (12,38), although in these studies, it may be the low transformation rate of the plasmid-based CRISPR tools. In our study, the loss of CRISPR activity may come from the burden of a two-plasmid system (17). In principle, both RP4 (as broad-host range conjugative plasmid) and the mobilized CRISPR plasmid should transfer to the new cell together. However, there can be circumstances where only the RP4 plasmid transfers successfully (20), diminishing its antibacterial activity. As such, a single conjugative plasmid system encoding both conjugative and CRISPR machineries may provide a solution and improve the efficiency of CRISPR-mediated killing (16–18,23,46). Although, it might require more work on the sequence optimization for broad host range and/or against wide targeted strains or selected genes.

In summary, our study shows that the success of CRISPR antimicrobial tools is highly dependent on community-specific ecological and evolutionary factors. Our study emphasizes that the parameters that govern the CRISPR-antibacterial in a complex community are still poorly understood, especially for a system that simulates real-life scenarios and more complex microbial environments. This work offers a useful insight into a possible outcome of using conjugative systems to deliver CRISPR antimicrobials for ESBL gene removal within complex communities and can be considered a useful reference for further study in natural systems, *e.g.*, gut microbiome, to improve the CRISPR antimicrobials to combat the rise in antibiotic resistance problem.

## Supplementary Tables

**Supplementary Table 1.**
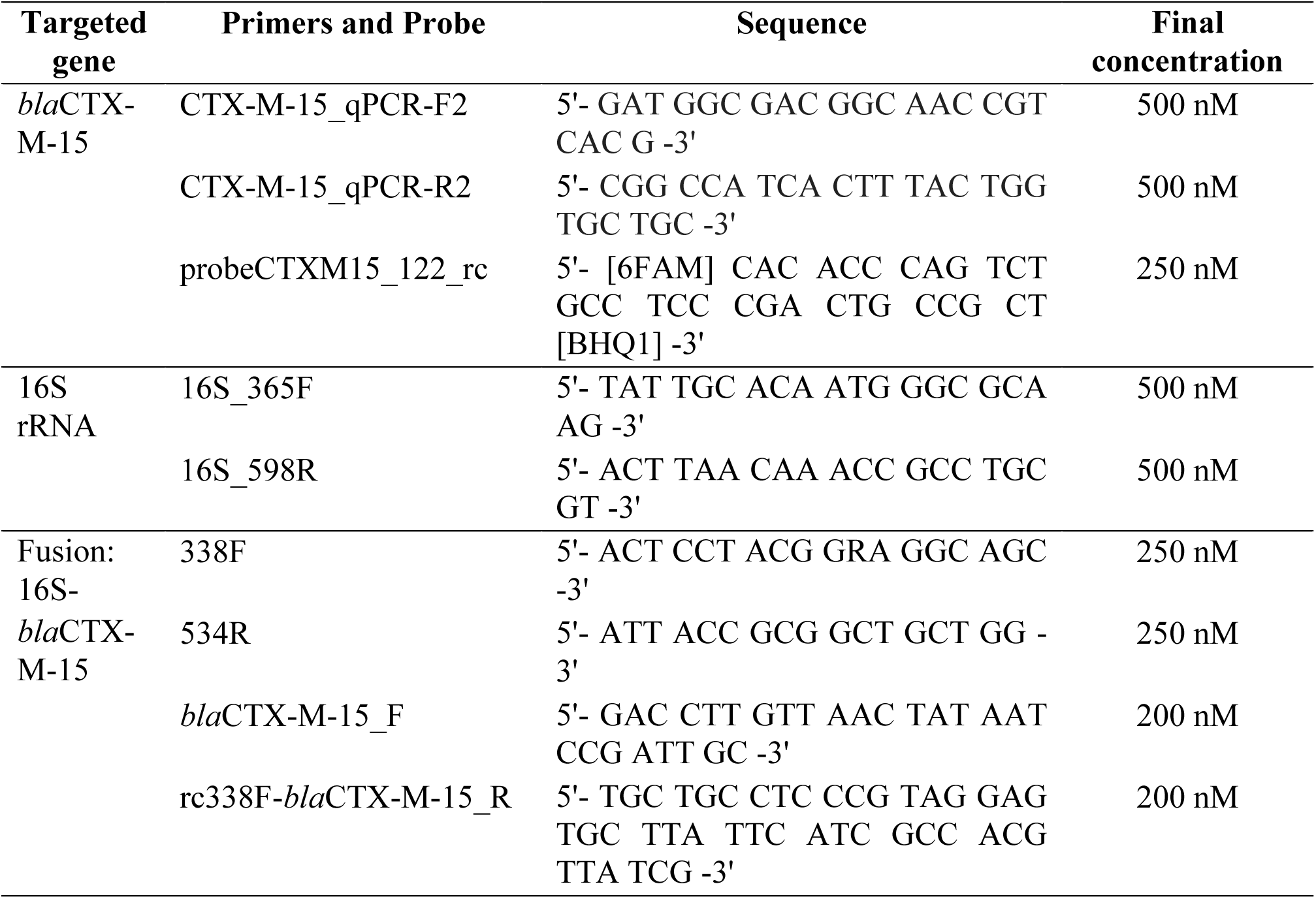
Primers and probe used in the experiment.

**Supplementary Table 2.** qPCR program for blaCTX-M-15 gene.

| Step | Temperature | Time |
| --- | --- | --- |
| 1. Initial denaturation | 95°C | 3 minutes |
| 2. Denaturation | 95°C | 15 seconds |
| 3. Primer annealing + Extension *plate read* | 65°C | 45 seconds |
| 4. Repeat step 2 to 3 for 40 times |  |  |

**Supplementary Table 3.** qPCR program for 16S.

| Step | Temperature | Time |
| --- | --- | --- |
| 1. Initial denaturation | 98°C | 3 minutes |
| 2. Denaturation | 98°C | 15 seconds |
| 3. Primer annealing + Extension *plate read* | 60°C | 1 minute 15 seconds |
| 4. Repeat step 2 to 3 for 40 times |  |  |
| 5. Melt curve *plate read* | 65 - 95°C, 0.5°C increment 5 seconds/step |  |

**Supplementary Table 4.** Program for fusion-PCR (*bla*CTX-M-15 & 16S).

| Step | Temperature | Time |
| --- | --- | --- |
| 1. Initial denaturation | 98°C | 30 seconds |
| 2. Denaturation | 98°C | 10 seconds |
| 3. Primer annealing | 55°C | 30 seconds |
| 4. Extention | 72°C | 15 seconds |
| 5. Repeat step 2 to 4 for 35 times |  |  |
| 6. Final extention | 72°C | 5 minutes |

**Supplementary Table 5.**
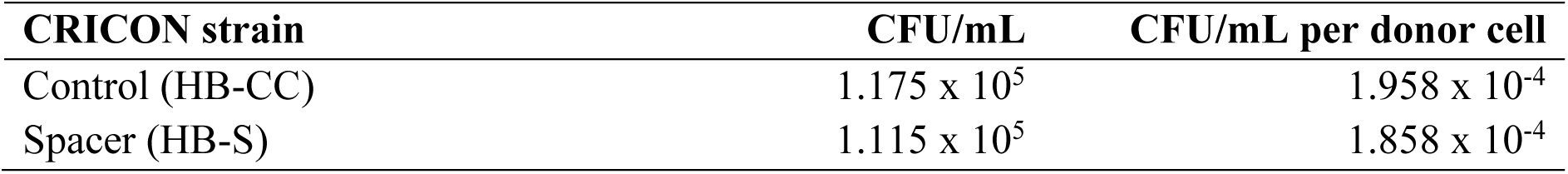
Average transfer rate of the CRICON strains into lab *E. coli* HMS174.

| CRICON strain | CFU/mL | CFU/mL per donor cell |
| --- | --- | --- |
| Control (HB-CC) | 1.175 x 10<sup>5</sup> | 1.958 x 10<sup>-4</sup> |
| Spacer (HB-S) | 1.115 x 10<sup>5</sup> | 1.858 x 10<sup>-4</sup> |

**Supplementary Table 6.** The ratio of initial CRICON donor: CRICON-ESBL transconjugant between the CRICON control (HB-CC) and spacer (HB-S) and the difference (Δ).

| | Ratio CRICON: transconjugant | | $\Delta$ Spacer – Control |
| --- | --- | --- | --- |
|  | Control (HB-CC) | Spacer (HB-S) |  |
| E4 | 0.00406 | 0 | -0.00406 |
| E5 | 0.00004 | 0 | -0.00004 |
| E6 | 0.03638 | 0 | -0.03638 |
| E7 | 0.00033 | 0 | -0.00033 |
| E8 | 0.00168 | 0.00003 | -0.00165 |
| E9 | 0.013985 | 0.00368 | -0.01031 |
| E10 | 0.0001 | 0 | -0.0001 |
| E11 | 0.00168 | 0.00022 | -0.00146 |
| E12 | 0.03623 | 0.00013 | -0.0361 |

**Supplementary Table 7.** Cross inhibition test between all the community strains used in the evolution microcosm experiment.

| Inhibiting strain \ Tested strain | HB-S | HB-CC | E4 | E6 | E8 | E11 | 4B24 | 4B41 | DSM 681<sup>rifS</sup> | HB101 |
| --- | --- | --- | --- | --- | --- | --- | --- | --- | --- | --- |
| HB-S |  | - | (+) | - | + | - | - | - | - | - |
| HB-CC | - |  | + | - | + | - | - | - | - | - |
| E4 (4B1) | - | - |  | - | - | - | - | - | - | - |
| E6 (12B1) | - | - | - |  | - | - | - | - | - | - |
| E8 (26B1) | - | - | - | - |  | - | - | - | - | - |
| E11 (77B1) | - | - | - | - | - |  | - | - | - | - |
| 4B24 (*Citrobacter freundii*) | - | - | - | - | - | - |  | - | - | - |
| 4B41 (*Klebsiella* sp.) | - | - | - | - | - | - | - |  | - | - |
| DSM681<sup>rifS</sup> | - | - | - | - | - | - | - | - |  | - |
| HB101 | - | - | (+) | - | + | - | - | - | - |  |

**NOTE:** + = positive inhibition, - = no inhibition, (+) = very thin clear zone surrounds the spot

## Supplementary Figures

**Supplementary Figure 1.**
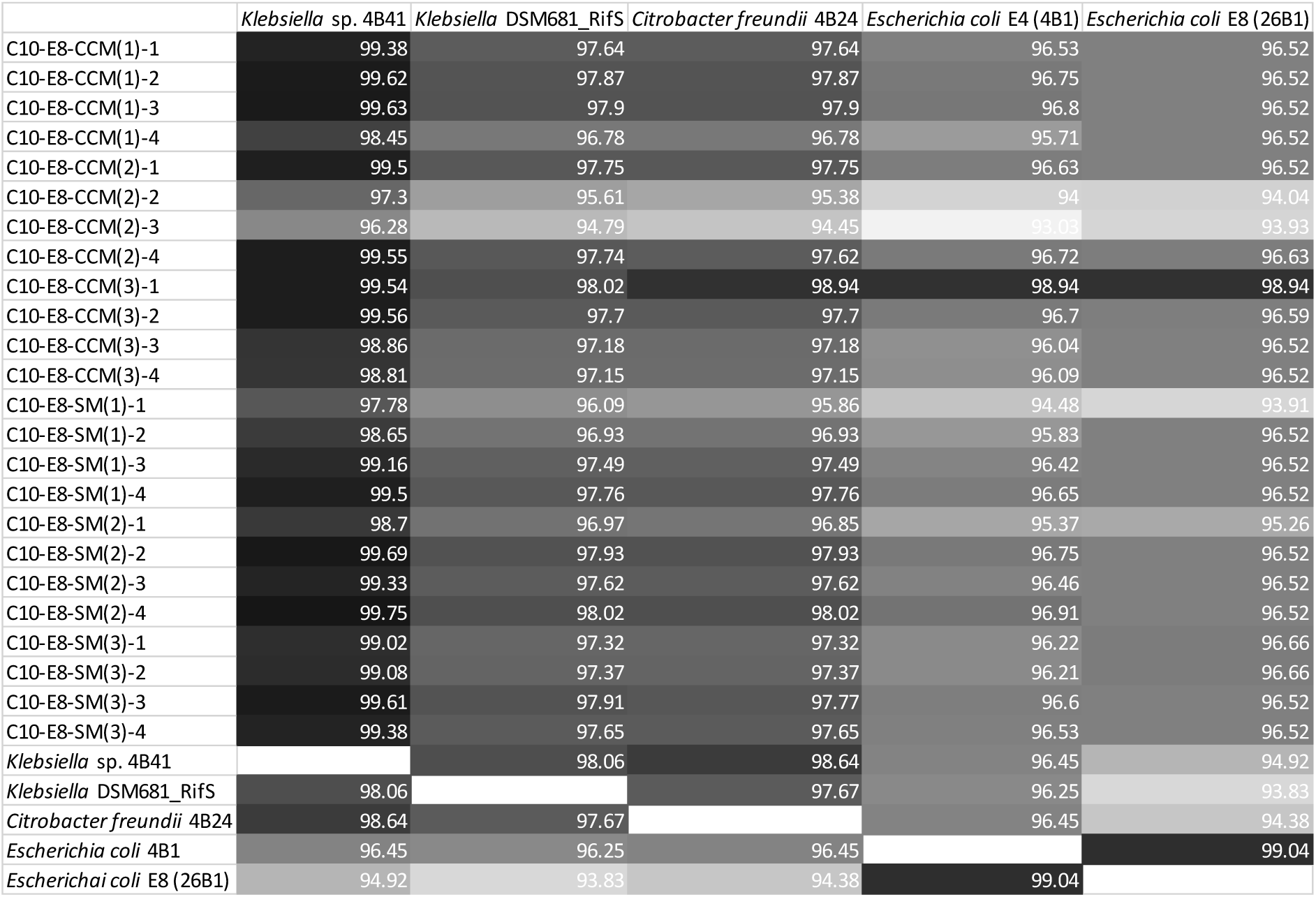
The alignment distance of E8 transconjugants with all the community strains based on 16S rRNA sequencing.

**Supplementary Figure 2.**
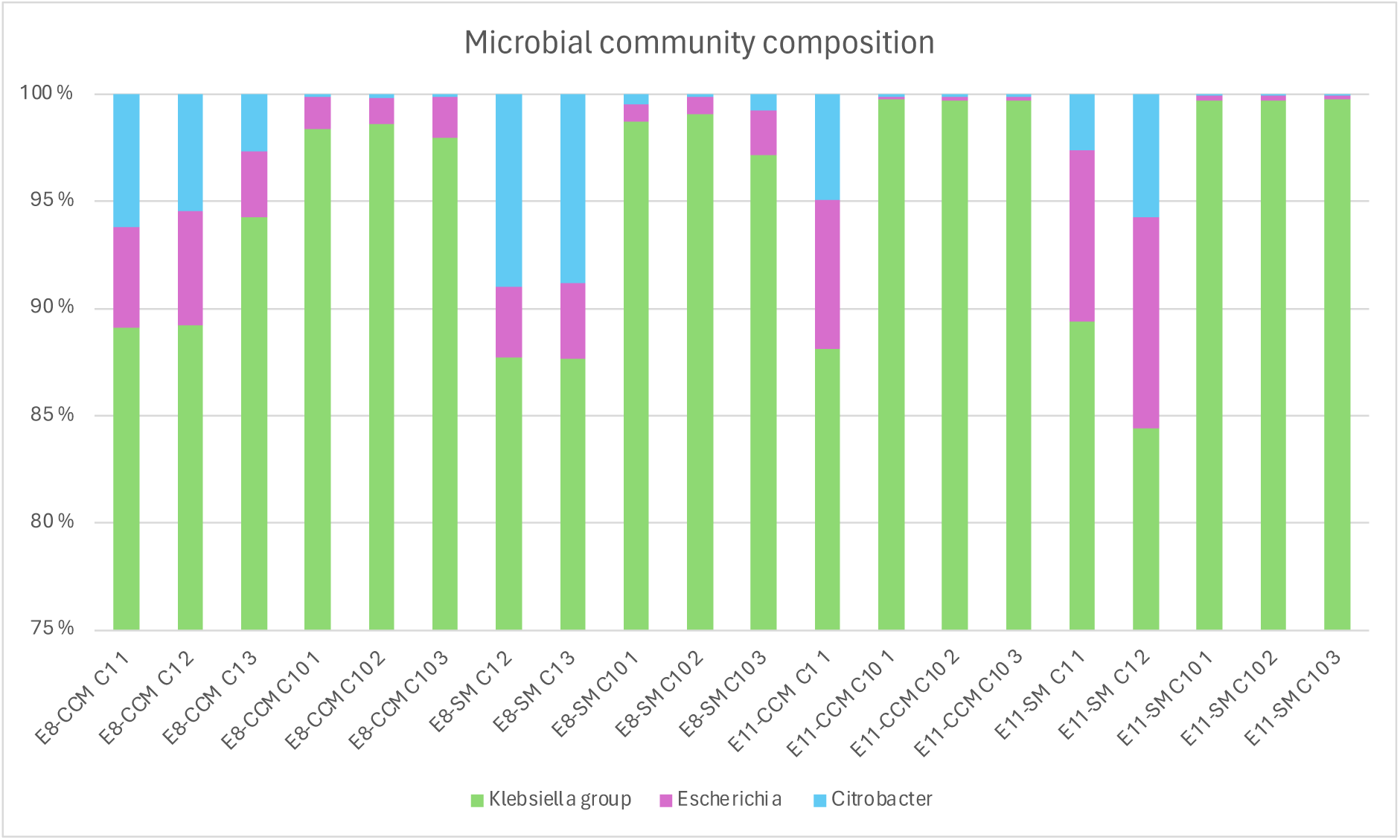
Community composition during the 10-day evolutionary experiment on day 1 (C1) and 10 (C10) in E8 and E11 communities. Community structure was determined by shallow shotgun metagenomic sequencing and taxonomic assignment based on read classification with kraken2 tool. CCM = CRICON control treatment with migration; SM = CRICON spacer (targeting *bla*CTX-M-15) treatment.

